# Identifying conceptual neural responses to symbolic numerals

**DOI:** 10.1101/2023.07.04.547627

**Authors:** Talia L. Retter, Lucas Eraßmy, Christine Schiltz

## Abstract

Neural processing of numerical concepts may be measured in humans automatically, without a related numerical task. However, the extent to which neural responses to symbolic numbers are due to physical stimulus confounds independently of conceptual representations remains unknown. Here, we targeted conceptual responses to parity (even *vs.* odd), using an electroencephalographic (EEG) frequency-tagging approach with a symmetry/asymmetry paradigm. Fifty second sequences of Arabic numerals (2–9) were presented at 7.5 Hz; odd and even numbers were alternated, so that differential responses to parity would be captured at 3.75 Hz (7.5 Hz/2). Parity responses were probed with four different stimulus sets, increasing in intra-numeral stimulus variability. Moreover, two control conditions were tested for each stimulus set, comprised of non-conceptual numeral alternations (strong control, for small inter-group physical differences: 2,3,6,7 *vs.* 4,5,8 and 9; weak control, for large physical differences: 2,4,5,7 *vs*. 3,6,8,9). Significant asymmetrical responses at 3.75 Hz were found over the occipitotemporal cortex to all conditions, thus even for arbitrary numeral groups. The weak control condition elicited the largest response in the stimulus set with the lowest level of variability (1 font). Only in the stimulus set with the highest level of variability (20 hand-drawn, colored exemplars per numeral) did the response to parity surpass both control conditions. These findings show that physical differences across small sets of Arabic numerals can strongly influence, and even account for, automatic brain responses. However, carefully designed control conditions and highly variable stimulus sets may be used towards identifying truly conceptual neural responses.

## Introduction

There is a longstanding interest in understanding the neural bases of numerical cognition. At a fundamental level, studies have addressed basic representations of numbers, such as numerosity and relative magnitude, with electroencephalography (EEG) and functional magnetic resonance imaging (fMRI; e.g., Dehaene, 1996; Fornaciai et al., 2017; Sokolowski et al., 2021). At a more abstract level, studies have also addressed the neural bases of operations (e.g., solving mathematical equations; Pauli et al., 1994; Jost, Henninghausen & Rosler, 2004; Park et al., 2012). Investigating the neural bases of numerical cognition may contribute to reconciling different theories of number processing, understanding brain organization more generally, and ultimately informing interventions in math learning.

An important issue in such studies is whether the observed neural responses are conceptual, or rather could be accounted for by physical stimulus features, such as variations in visual properties. Visual stimulus confounds (such as dot size, density, total area, cumulative perimeter, total luminance, dot shape, array shape, etc.) have long been known to influence behavioral responses to non-symbolic representations of numbers, in particular dot arrays (see Dehaene, Izard, & Piazza, 2005; Gebuis, Kadosh & Gevers, 2016; Cheng et al., 2023). As in behavioral studies, neural responses intended to reflect cognitive processing for non-symbolic dot representations, such as numerosity, are typically either contrasted to responses to selected visual stimulus properties (Soltesz & Szucs, 2014; Park, 2018; Van Rinsveld et al., 2020) or attempted to be isolated by orthogonally varying multiple visual stimulus features (Hyde & Spelke, 2011; Libertus, Brannon & Woldorff, 2011; Guillaume et al., 2018; Marlair et al., 2021; but see Georges, Guillaume & Schiltz, 2020).

However, the issue of whether neural responses are affected by physical stimulus features is outstanding in the case of symbolic representations of numbers. One reason for this may be that it is difficult to identify physical confounds, such as line curvature, closure, or spatial frequency, that may influence responses as measured with different techniques. However, symbolic representations of numbers use exemplars that are highly distinctive in their physical features, enabling easy and rapid identification (e.g., for rapid identification of western Arabic numerals: Starrfelt & Behrmann, 2011). This may account for the possibility for above-chance decoding of the identity of Arabic numerals with fMRI (Eger et al., 2009; Bluthe, De Smet & Op de Beek, 2014) and EEG (Appelhoff, Hertwig & Spitzer, 2022). Physical differences across numerals may also strongly influence behavioral responses: physical differences from the number 5 predict the numerical distance effect (since Moyer & Landauer, 1967) across Arabic numerals from 1 to 9 much better than the quantity of numerical distance (Cohen, 2009).

In addition to being highly distinctive, the limited stimulus variability of symbolic representations of numbers may contribute to the impact of their physical attributes. Arabic numerals rely on very few basic exemplars, i.e., ten numerals from 0 to 9. Additionally, numerals are typically presented with approximate homogeneity in most aspects, such as size, line thickness, and color. Most studies using Arabic numerals have used a single font type and point size, in black or white (e.g., Dehaene, 1996; Fabre & Lemaire, 2005; Libertus & Brannon, 2007; but for more variation in the context of adaptation, see Vogel, Goffin & Ansari, 2015; Finke et al., 2021). Stimulus variability is important, in that it relates to the consistency, and thus impact, of physical stimulus confounds on responses: increasing stimulus variability has been identified as a way to decrease stimulus confounds (e.g., with images: Thorpe et al., 1996; Foldiak et al., 2004; Rossion et al., 2015; with dots: Georges, Guillaume & Schiltz, 2020; with written words and/or numbers: Cohen, 2009; Lochy & Schiltz, 2019; Volfart et al., 2020; Marlair, Crollen & Lochy, 2022). Taken together, the limited variability within symbolic number exemplars, and the high variability across number exemplars, leaves responses highly susceptible to stimulus confounds.

In many previous studies using Arabic numerals, multiple number formats or modalities, or many exemplars of mathematical equations, have been used instead of controlling for numerals’ visual confounds (e.g., Pauli et al., 1994; Dehaene, 1996; Liu et al., 2018; van Hoogmoed & Kroesbergen, 2018). The issue of stimulus confounds for symbolic representations of numbers is most pertinent when attempting to distinguish conceptual neural responses, such as for parity or magnitude, with restricted stimulus sets (e.g., Hsu & Szucs, 2012; Guillaume et al., 2020; Sokolowski et al., 2021). Take the example of *parity*, a conceptual distinction that divides numbers into the categories of even and odd. These categories are defined by the ending of numbers in one of five digits (0,2,4,6,8 for even; 1,3,5,7,9 for odd); given the extreme differences in shape features across Arabic numerals, the even set of numerals is more curved, closed, and symmetrical than the odd set. Therefore, neural responses distinguishing even and odd numerals (Fabre & Lemaire, 2005; Guillaume et al., 2020) might reflect a conceptual distinction of parity categories, but could also reflect the visual system’s sensitivity to physical differences in numeral shape features across small groups of even and odd numbers.

Here, we targeted the conceptual numerical processing of parity, using EEG frequency-tagging to differentiate neural responses to a group of even (2,4,6,8) and a group of odd (3,5,7,9) Arabic numerals. Importantly, we tested two control conditions of non-conceptual numeral groups, designed to differ on the extent of their physical stimulus differences across groups (*strong control*, for small physical differences: 2,3,6,7 *vs.* 4,5,8,9; and *weak control*, for large physical differences: 2,4,5,7 *vs.* 3,6,8,9; **Fig. 1**). Moreover, we presented four different stimulus sets orthogonally, differing in their amount of variability: 1) *1 font* (lowest variability); 2) *10 fonts*; 3) *10 mixed* fonts/numeral drawings; and 4) *20 drawn* diverse, colored numerals (highest variability). We explored whether conceptual responses to parity could be distinguished from those to physical differences of non-conceptual number groups, quantitatively and qualitatively, and whether this depended on stimulus set variability. We predicted that the weak control would yield larger response amplitudes than the strong control, particularly with lower levels of stimulus set variability. Most importantly, we predicted that the response to parity would be larger than both control responses only with higher levels of stimulus set variability.

**Figure 1.**
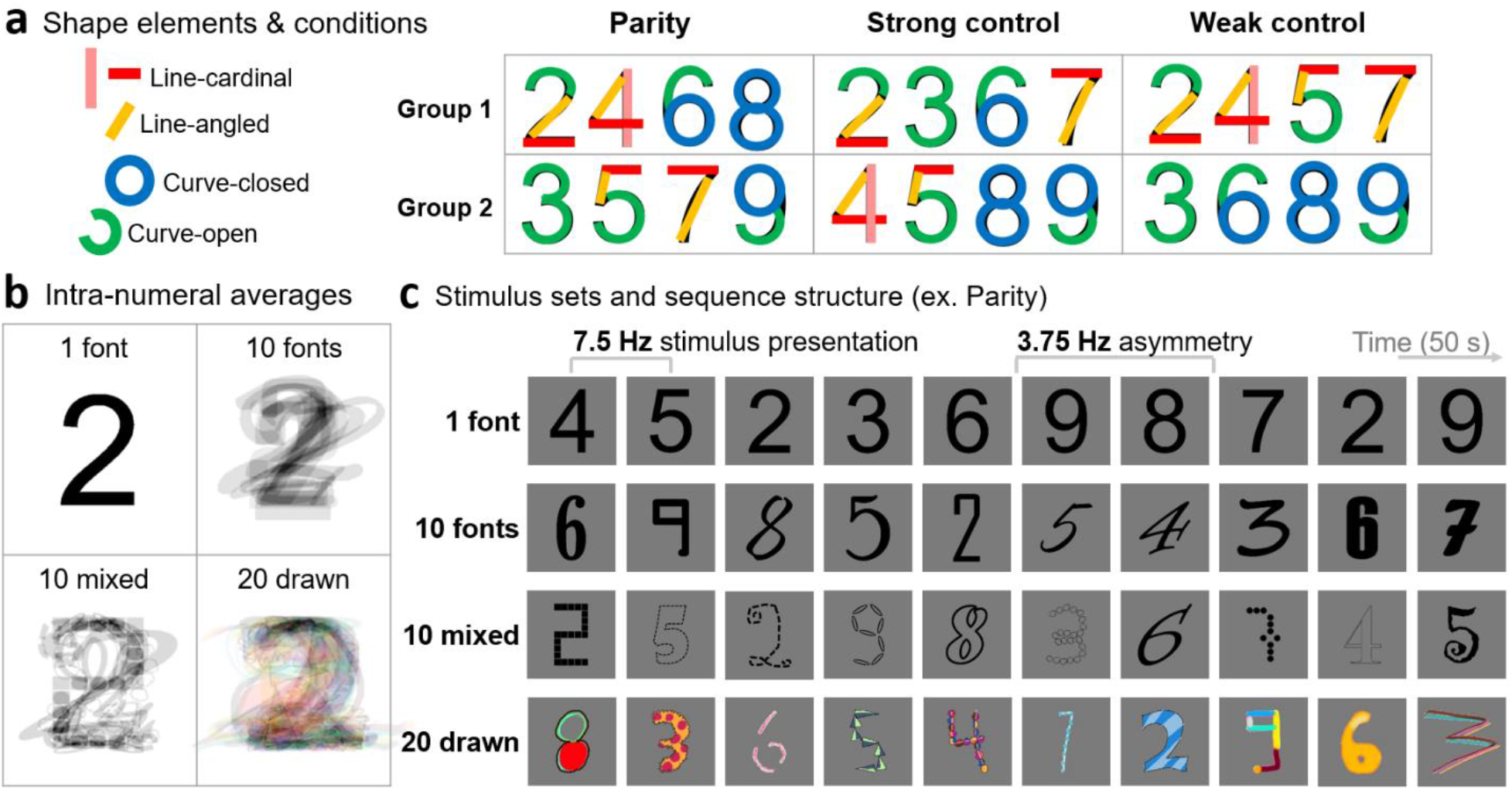
Experimental design. ***a)** Three experimental conditions were each defined by two groups of four Arabic numerals. In the Parity condition, group 1 consisted of even numerals, and group 2 of odd numerals. In the control conditions, numerals were assigned non-conceptually to groups in consideration of the amount of shape differences across groups (small inter-group differences in the Strong control; large in the Weak control). **b)** Intra-numeral averages, with the numeral 2 as an example, demonstrating the variability of each of the stimulus sets. **c)** Stimuli were presented in 50 s sequences at 7.5 Hz (133 ms cycles), alternating between numerals of the two groups of each condition, leading to a 3.75 Hz asymmetry frequency tag.*

## Methods

### Participants

Participants were 15 healthy, neuro-typical human adults (age range: 18 – 27 years; age mean = 23.4 years; 2 males; 13 females). All 15 participants reported to have normal or corrected-to-normal vision, and all but two to be right-handed. The sample size approximately matched that of previous frequency-tagging studies reporting strong effects, e.g., using a symmetry/asymmetry paradigm: Retter & Rossion, 2016a; in numerical cognition: Park, 2018). They were recruited from the University of Luxembourg community via educational and social media. Each participant was tested individually, following signed, informed consent, with procedures approved by the Ethical Review Panel of the University of Luxembourg (ERP 20-057), and consistent with the Code of Ethics of the World Medical Association (Declaration of Helsinki).

### Stimuli and conditions

The stimuli were the eight Arabic numerals ranging from 2-9. The digit 0 was excluded as it is not always readily identified as even, and 1 was arbitrarily excluded so that two groups of four numerals could be defined. This also corresponds with the numerals that have been used in many previous EEG studies (Fabre & Lemaire, 2005; as operands: Pauli et al., 1994; Niedeggen, Rosler & Jost, 1997; Rosler & Jost, 1999; Jost, Henninghausen & Rosler, 2004; Zhou et al., 2006; Domahs et al., 2007). Three experimental conditions allowed the comparison of *Parity* with two controls, *Strong control* and *Weak control*. In the Parity condition, the even integers 2,4,6, and 8 defined group 1, and the odd integers 3,5,7, and 9 defined group 2. In the Strong control: group 1 consisted of 2,3,6, and 7, and group 2 consisted of 4,5,8 and 9. In the Weak control: group 1 consisted of 2,4,5 and 7, and group 2 consisted of 3,6,8, and 9 (**Fig. 1a**).

The control conditions were designed to provide two relative measures of non-conceptual, visual differences across numeral groups: these groups did not differ in parity, each containing 2 even and 2 odd numbers. The groups were also approximately matched for magnitude: Parity group 1 *vs.* 2 means: 5 *vs.* 6; Strong control: 4.5 *vs.* 6.5; Weak control: 4.5 *vs.* 6.5. In the Strong control, intended to have lower visual differences across numeral groups, the basic shape features of straight (along the cardinal axes or angled) *vs.* curved contours, open *vs.* closed elements, and symmetry (e.g., Triesman, 1986; McCloskey & Schubert, 2014; for features in letter perception: Fiest et al., 2008; Grainger, Rey & Dufau, 2008) of the Arial font were approximately balanced across stimulus groups by visual inspection. In the Weak control, the curviest, the most closed numeral shapes were sorted into a single group (3,6,8 and 9). None of these integers contained a straight line, and three contained at least one closed element; while all of the numerals in the other group (2,4,5 and 7) contained at least one straight line, and only one of which contained one closed element. Two numerals also had horizontal symmetry in group 2 of the Weak control (3 and 8), while none did in group 1.

The amount of visual differences across groups for each of the conditions was also measured quantitatively, for the Arial font, in terms of non-background image space and spatial frequency (using the statistical toolbox of Bainbridge & Oliva, 2015), as well as parametric complexity (Attneave & Arnoult, 1956). The amount of physical similarity of all stimulus pairs within each group was also measured (using the method of Cohen, 2009). For all these properties, the differences across numeral groups for Parity and the Strong control were similar, tending to be somewhat smaller for Parity; the differences across numeral groups for the Weak control were considerably the largest (**Table S1**). The Weak control was thus tested as a dramatic example of the possible contribution of physical stimulus confounds, rather than a control intended to match the confounds of the Parity condition *per se*.

### Stimulus sets

There were four different stimulus sets, designed to differ in their intra-numeral variability (**Fig. 1b**). In the *1 font* stimulus set, only the Arial font was used, defining one exemplar of each numeral. In the *10 font* set, the variable fonts 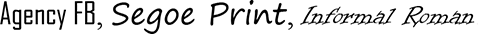, Poor Richard, 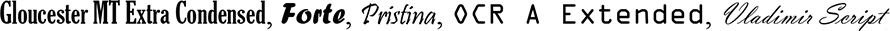, and 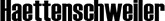 were used, so that there were 10 exemplars of each numeral. Note that including variability by changes in font has been applied in previous studies with Arabic numerals (e.g., Cohen, 2009; Vogel, Goffin & Ansari, 2015; with EEG: Lochy & Schiltz, 2019; Guillaume et al., 2020; Finke et al., 2021; Marlair, Crollen & Lochy, 2022). The *10 mixed* set was also comprised of 10 exemplars of each numeral, but designed to include even more variability across exemplars that included both fonts and drawings. There were five fonts used: two in plain type (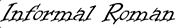 and 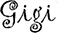), two in outline only, i.e., with a transparent interior (Modern No. 20 and 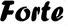), and one in a dotted outline only 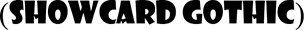. Additionally, there were five computer-drawn exemplars, comprised of: black squares in a grid organization, black dots in a grid with gentle edges, outlined cloud shapes, outlined ovals, and a free-form dotted line. Finally, the *20 drawn* set was comprised of 20 exemplars of each numeral, hand-drawn in color on a tablet (details below). Throughout, exemplar style (e.g., blue diagonal stripes with a thick angular shape in *20 drawn*) was applied to each of the numerals, to decrease inter-numeral variability.

In stimulus generation, the font and computer-drawn stimuli were generated in PowerPoint (Microsoft, USA). The hand-drawn stimuli were generated with a stylus (Smart Pen, Dongguan Ruibi Touch Technology Co., Ltd., China) on a tablet (iPad, 10.2 inch screen, Apple, USA). The numbers were extracted as PNGs on a white background, before being auto-cropped to the borders of the number edges in IrfanView 64 4.53 (https://www.irfanview.net/). The stimuli were seized to a height of 360 pixels, based on the Arial font only for all font stimuli; variability in height across fonts of the same text size was retained. Finally, the image canvas was expanded from the center to a common size of 500×500 pixels, and the background was set to a mid-level gray (123/255). All four stimulus sets will be made freely available online upon manuscript acceptance (including the numerals 0 and 1 for the *20 drawn* stimulus set): for some examples, see **Fig. 1**.

### Paradigm and procedure

To record neural responses to parity, electroencephalogram (EEG) was recorded with a frequency-tagging approach (also referred to as “steady-state visual-evoked potentials” or “fast periodic visual stimulation”; Regan, 1989; Rossion, 2014; Norcia et al., 2015). Stimuli were presented at a periodic rate, 7.5 images a second (7.5 Hz), and the recorded EEG was transformed into the frequency domain, in which responses to stimulus presentation are specifically tagged at 7.5 Hz and its harmonics (**Fig. 2**). We applied a symmetry/asymmetry design (Tyler & Kaitz, 1977; Victor & Zemon, 1985; for more recent examples, Ales et al., 2012; Retter & Rossion, 2016a), in which the two groups of images (e.g., the even and odd numerals for the parity condition) within each stimulus set were alternated, leading to a 3.75 Hz (i.e., 7.5 Hz/2) tag of asymmetrical responses to the contrasted integer groups. In the parity conditions, these responses are predicted to reflect conceptual differences in the neural responses to even and odd numbers, although physical differences are also expected to contribute, as measured specifically in the control conditions.

**Figure 2.**
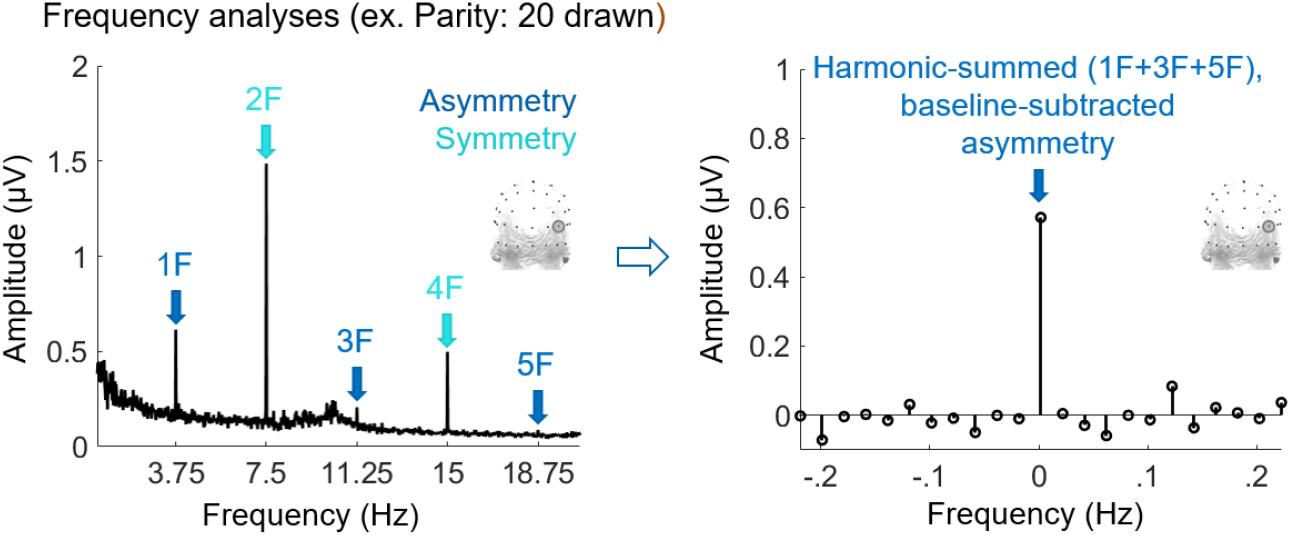
Harmonic-summation and baseline-subtraction. *Group-level experimental data from the parity condition, with the 20 drawn stimulus set, are given as an example (electrode PO8) of the original frequency-domain amplitude spectrum, and the measurement of asymmetry (and symmetry) responses through harmonic-summation and baseline-subtraction. In the right panel, 0 Hz is the target frequency across harmonics, surrounded by a relative frequency range.*

Stimuli were presented as a 62.5% squarewave modulation at 7.5 Hz, i.e., appearing for 83 ms followed by a 50 ms inter-stimulus interval. Stimuli were presented through Java (Oracle, USA), with a Dell S2419HGF (USA) monitor with a screen size of about 53×30 cm and a refresh rate of 120 Hz, connected to a Dell (USA) PC with a Geforce 1050 (Nvidia, USA) graphics card. Participants had a 1 m viewing distance, such that stimuli subtended about 1.9° of vertical visual angle on average. To reduce exact image repetitions, at each presentation cycle the size of the stimulus was varied randomly between 80-120% of the original size in 10% increments (Dzhelyova & Rossion, 2014). Throughout stimulation, participants were asked to fixate on a centrally presented dark gray fixation cross, and were instructed to press on the space bar each time the cross briefly (250 ms) changed to white, which occurred 6 times per sequence at random intervals, with a minimum of 750 ms between changes (color task : Park, 2018; Guillaume et al., 2018; Van Rinsveld et al., 2020; passive viewing: Libertus, Brannon & Woldorff, 2011; see the Supplemental material for the behavioral results).

Each stimulation sequence lasted for 50 s, throughout which stimuli from the two groups of each condition were alternated, leading to an asymmetry frequency tag at 3.75 Hz. In the full trial structure, designed to maintain fixation and avoid artifacts at the beginning and end of stimulation, there was: 1) 1-2 s of the fixation cross alone; 2) 2 s of gradual stimulus fade-in; 3) the 50 s stimulation sequence; 4) 2 s of gradual stimulus fade-out; and 5) 1-2 s of the fixation cross. For each condition there were three trial repetitions, leading to a total of 2.5 minutes of the stimulation sequence. This time encompassed 562 stimulus asymmetry presentations per condition. In total, including rest times between trials, the experimental recording session lasted about 40 minutes.

### EEG acquisition

EEG was acquired with a BioSemi, ActiveTwo system, comprised of 68 active recording electrodes in the standard 10/20 locations; the standard 64, plus PO9, I1, I2, and PO10 (BioSemi B.V., Amsterdam, Netherlands). Electrode offsets were held below 40 mV, relative to the system’s common-mode-sense and driven-right-leg reference loop. The EEG was digitized at a sampling rate of 512 Hz.

### Analysis

Analysis was conducted with LetsWave 6 (https://www.letswave.org/), running over Matlab R2019b (MathWorks, USA), largely in accordance to previous symmetry/asymmetry designs (Retter & Rossion, 2016a; 2017). Briefly, and noting any differences: data were pre-processed, including a fourth-order Butterworth band-pass filter at 0.1-80 Hz, and coarsely segmented 1 s before and after each trial. Muscular artifacts related to eye blinks were corrected by removing a single ICA component, for participants blinking more than 0.2 times per s during the stimulation sequences (three participants; overall M = 0.14 blinks/s; SD = 0.14 blinks/s). Noisy channels, containing multiple deflections over ± 100 µV, were corrected for by linear interpolation with neighboring channels (M = 1.4 channels per participant; range = 0-3). Data were re-referenced to the average of all EEG channels, and the stimulation sequences were precisely cropped to an integer number of 3.75 Hz cycles (187 cycles per sequence = 49.87 s = 25,532 sampling bins). Sequence repetitions were averaged together by condition, before a fast Fourier transform was used to represent the data as frequency-domain amplitude spectra (**Fig. 2**), and phase spectra.

The harmonics of 3.75 Hz in the amplitude spectrum were segmented, together with a range of neighboring frequency “noise” bins. A baseline was defined by a range of about 0.4 Hz surrounding noise (20 bins; for the baseline subtraction, the minimum and maximum bin were excluded). For response amplitude measurement, a baseline subtraction was applied; for significance assessment of individual harmonics, a Z-score was computed relative to this baseline. The harmonics-of-interest were defined by those consecutively surpassing a threshold of p < .01, Z > 2.32, 1-tailed, over the average of all EEG channels, in at least one condition (Retter, Rossion & Schiltz, 2021). This led to three odd, asymmetry harmonics: 3.75, 11.25, and 18.75 Hz; and six even, symmetry harmonics: 7.5, 15, 22.5, 30, 37.5, and 45 Hz. These harmonics were then summed for further analyses, separately for the asymmetry and symmetry responses (**Fig. 2**; Retter & Rossion, 2016b; Retter, Rossion & Schiltz, 2021).

The region-of-interest (ROI) was defined as a bilateral occipitotemporal region, containing eight channels (PO8; PO10; O2; I2; and corresponding left channels; drawn in **Fig. 3**). This ROI was originally based on previous studies using numeral stimuli, in which responses were maximal over occipitotemporal regions, with responses typically peaking around PO8 (see Fig. 3 of Libertus, Woldorff & Brannon, 2007; Fig. 2 of Liu et al., 2018; Fig. 4 of Akbari et al., 2022; Pinel et al., 2001; Plodowski et al., 2003; Guillaume et al., 2020; Marinova et al., 2021), although sometimes labeled as an (occipito)parietal region. Post-hoc, it was verified that this ROI appropriately captured the amplitude distribution of the response over the scalp: it contained on average 76% of the top eight channels across conditions and stimulus sets for the harmonic-summed, baseline-subtracted asymmetry responses of interest.

**Figure 3.**
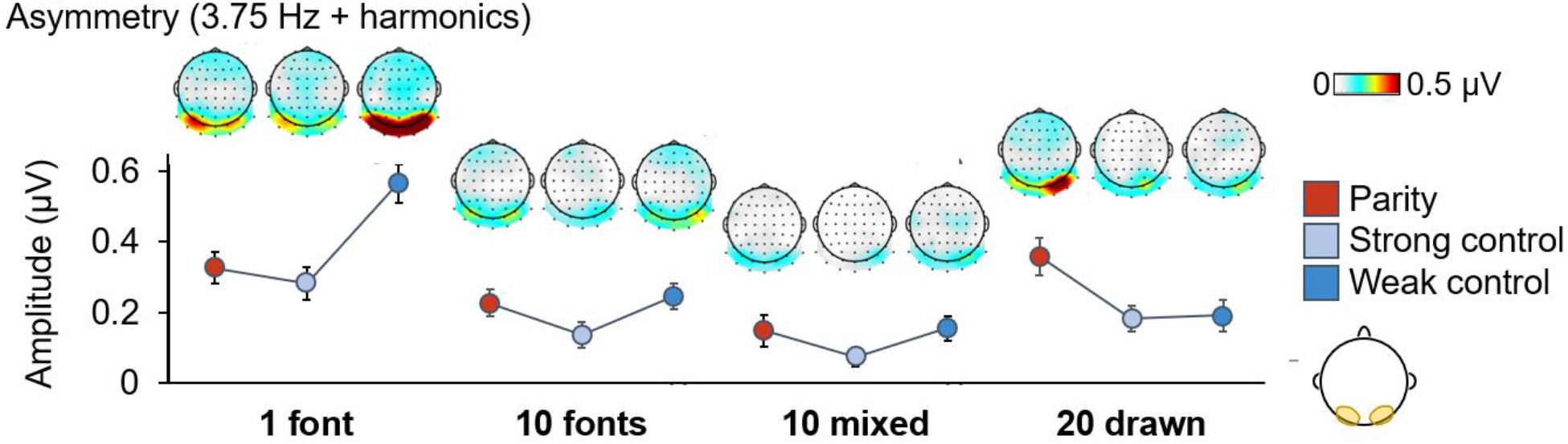
Asymmetry responses at 3.75 Hz and harmonics: group-level amplitudes and scalp topographies. *Amplitude is quantified over the occipito-temporal ROI, indicated on the scalp plot in the bottom right, harmonic-summed and baseline-subtracted. Error bars indicate ± 1 SE of the mean.*

**Figure 4.**
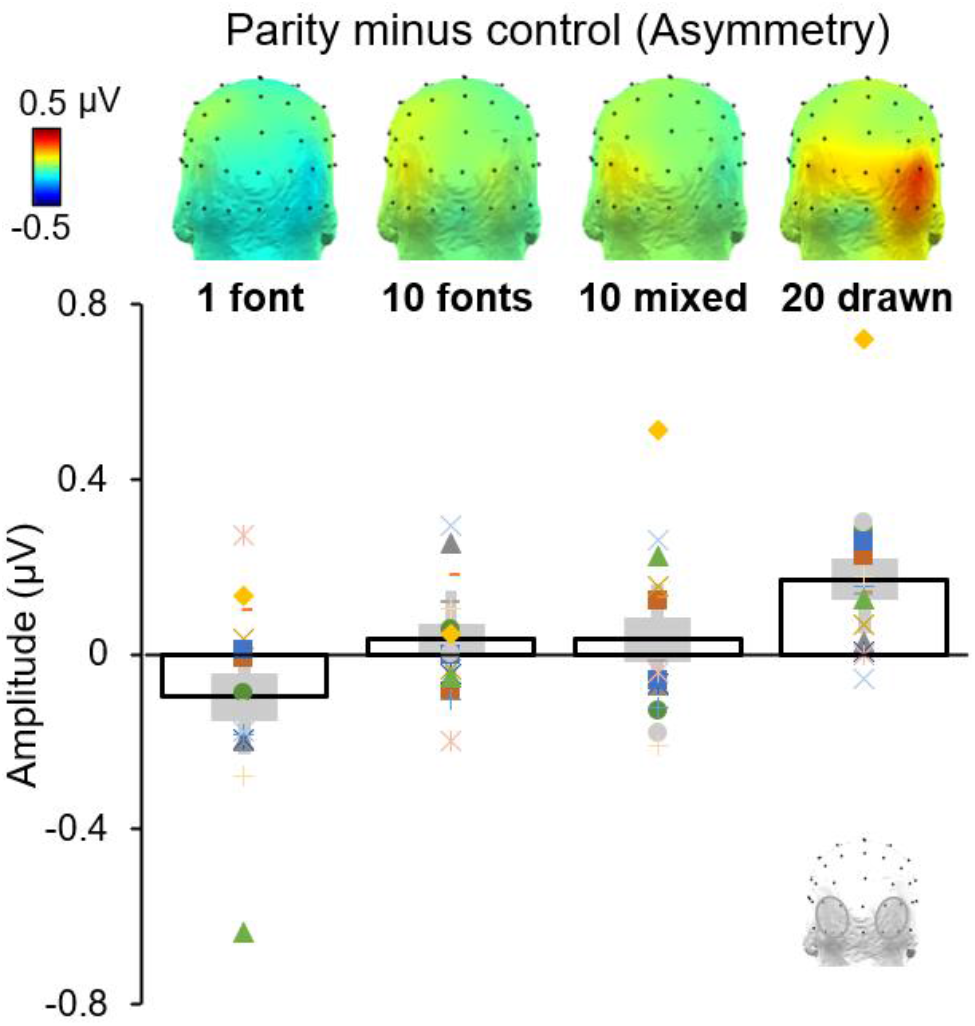
Asymmetry response amplitude differences for parity minus the average of the two control conditions. *Amplitudes from 3.75 Hz are harmonic-summed and baseline-subtracted. Above: back-of-the-head, 3D scalp topographies. Below: the corresponding response amplitudes over the occipito-temporal ROI, as drawn in the bottom right. Mean data is plotted with a black outline bar plot, with ± 1 SE of the mean indicated by a filled gray rectangle. Individual participant data are represented with dot markers, with each individuals’ marker shape and color held constant across stimulus sets.*

To determine significance of asymmetry responses overall, a Z-score was assessed on the summed-harmonic responses over the occipito-temporal ROI (Retter, Rossion & Schiltz, 2021). A repeated-measured analysis-of-variance (ANOVA) was applied, with within-participants factors of *Condition* (3 levels: Parity; Strong control; Weak control), and *Stimulus Set* (4 levels: 1 font; 10 fonts; 10 mixed; 20 drawn). Follow-up one-way ANOVAs were then performed separately for each stimulus set, with the factor of *Condition*. Further, follow-up, paired-samples t-tests were performed to compare the conditions within stimulus sets in a pairwise manner (3 comparisons, Bonferroni-corrected, 2-tailed).

## Results

Frequency-domain amplitude spectra were analyzed through harmonic-summation and baseline-subtraction (**Fig. 2**). The main responses of interest, at 3.75 Hz and its specific harmonics, tagged asymmetrical responses, reflecting differences in the responses to the alternated numeral groups. The results showed asymmetry response amplitudes peaking over the occipitotemporal cortex that varied across conditions and stimulus sets (**Fig. 3**).

An asymmetry response to Parity (2,4,6,8 *vs.* 3,5,7,9) was present significantly for all stimulus sets; however, the asymmetry responses were also significant in all the control conditions, all Z’s > 2.32, p’s <.01 (for exact values, see **Table S2**).

There were significant main effects of *Condition,* F_2,28_ = 8.16, p = .002, ɳ_p_^2^ = 0.37, and *Stimulus Set,* F_3,42_ = 28.5, p < .001, ɳ_p_^2^ = 0.67, for the asymmetry responses. However, these main effects were qualified by a significant interaction, F_6,84_ = 6.04, p < .001, ɳ_p_^2^ = 0.30. Follow-up analyses within stimulus sets revealed a significant effect of *Condition* only for 1 font, F_2,28_ = 12.3, p < .001, ɳ_p_^2^ = 0.47, and 20 drawn, F_2,28_ = 8.10, p = .002, ɳ_p_^2^ = 0.37 (10 fonts: F2 = 2.64, p = .089, ɳ_p_^2^ = 0.16; 10 mixed: F2 = 1.62, p = .22, ɳ_p_^2^ = 0.10).

For 1 font, this was driven by the largest response for the Weak control (p’s < .003); the difference between Parity and the Strong control was not significant, t_14_ = 0.66, p = .16, d = .25. For 20 drawn, the effect of *Condition* was driven by the largest response for Parity (p’s < .034); there was not a significant difference between the two control conditions, t_14_ = 0.21, p = 2.5, d = .056. That is, while Parity responses were generally higher than those to the Strong control across stimulus sets, only the 20 drawn stimulus set produced significantly larger asymmetry responses than both the control conditions (**Fig. 4**).

The scalp topography of Parity asymmetry responses appeared similar to those of the control conditions within each stimulus set, peaking over occipito-temporal channels; but differences in lateralization were apparent across stimulus sets (**Fig. 3**). In contrast, the topography of symmetry responses appeared relatively more consistent across stimulus sets, and did not appear to differ by condition (**Fig. S1**). Differences in the asymmetry responses to Parity *vs.* control conditions within the 20 drawn stimulus set were more distinctively observed in the variability across individual participants, which appeared to be decreased relative to the controls in terms of both topography and phase (**Fig. S2**).

Finally, the symmetry responses at 7.5 Hz and its harmonics, tagging symmetrical responses to visual stimulus presentation, were examined. These responses did not considerably differ across conditions for any of the stimulus sets (**Fig. S1**). There was only a significant main effect of *Stimulus Set,* F_3,42_ = 5.46, p = .003, ɳ ^2^ = 0.28 (all other *F’s* < 0.55, p’s > .77). This reflected small response amplitude differences overall across stimulus sets, as follows in decreasing order: 10 fonts, 1 font, 20 drawn, 10 mixed (respectively, M = 2.14 µV, SE = 0.13 µV; M = 2.00 µV, SE = 0.15 µV; M = 1.87 µV, SE = 0.093 µV; M = 1.78 µV, SE = 0.096 µV).

## Discussion

To record responses to a numerical concept, parity, we used an EEG frequency-tagging paradigm, in which even (2,4,6,8) *vs.* odd (3,5,7,9) symbolic, Arabic numerals were contrasted at 3.75 Hz. Responses at 3.75 Hz thus reflected differences in the conceptual responses to even and odd numerals – as well as to any elicited visual response differences between the groups of numerals. To isolate responses to physical stimulus differences, we tested two control conditions in which groups of non-conceptual numerals were alternated at 3.75 Hz (Strong control, targeting small physical differences across groups: 2,3,6,7 *vs.* 4,5,8,9; Weak control, targeting extremely large physical differences: 2,4,5,7 *vs.* 3,6,8,9). To further investigate the influence of visual attributes of the EEG responses, we compared four stimulus sets, differing in their variability (from 1 font, consisting of one Arial font exemplar per numeral, up to 20 drawn, consisting of 20 hand-drawn, colored exemplars per numeral). We investigated whether Parity would give larger and/or qualitatively different 3.75 Hz asymmetry responses than the control conditions, and to what extent this was supported by increasing stimulus variability.

### Identifying conceptual responses to parity with variable stimuli

The asymmetry response at 3.75 Hz to Parity over the occipitotemporal cortex was significantly larger than to the two control conditions only for the 20 drawn stimulus set, and appeared larger than to the Strong control for the 10 font and 10 mixed stimulus sets. The asymmetry response to Parity was not significantly larger than either control condition for the 1 font stimulus set. A higher response amplitude to Parity than the control(s), particularly as stimulus set variability increases, is in line with our prediction that conceptual responses to parity may be evidenced beyond (variable) physical stimulus influences. An automatic access to parity would be consistent with some previous studies (e.g., Krueger & Hallford, 1984; Reynvoet et al., 2002; with EEG: Fabre & Lemaire, 2005; Guillaume et al., 2020), and with the status of parity as a fundamental aspect of number knowledge (Ifrah, 2000). However, the interpretation of capturing automatic, conceptual responses to parity, building upon the persistent response amplitudes to physical stimulus differences as evidenced in the control condition(s), must be proposed tentatively: physical stimulus differences can equal or surpass the amplitude of potential conceptual response differences to even and odd numerals, and there was a lack of qualitative response specificity for Parity (as addressed in a following section).

For the 1 font stimulus set, despite changes in presentation size, the Parity asymmetry response was not substantially larger than the Strong control, and was much smaller than the Weak control. With little stimulus variability, the visual response amplitude generated by physical stimulus attributes might be considerable enough to exceed a conceptual response component. With moderate variability in the 10 fonts and 10 mixed stimulus sets, the Parity asymmetry responses appeared higher than the Strong control (10 fonts: Parity was 1.7 times larger than the Strong Control; 10 mixed: 2.0 times). For the most variable, 20 drawn stimulus set, the Parity response was two times larger than the Strong control, and interestingly, 1.9 times larger than the Weak control. That is, relative to the other stimulus sets, the response to the Weak control appeared selectively decreased (**Fig. 3**).

One interpretation is that the low-to mid-level visual response differences elicited by the numeral groups were more equated across conditions for this highly variable stimulus set, leading to the least ambiguous identification of conceptual Parity responses. This stimulus set created the highest amount of diversity within numeral exemplars (20 colored, hand-drawn per numeral), while decreasing the variability across numerals (each schema was applied to all numerals). Increasing variability intra-numerally and decreasing variability inter-numerally might decrease the relative importance of the latter, leading to a smaller impact of visual differences across numeral sets (e.g., Thorpe et al., 1996; Foldiak et al., 2004; Rossion et al., 2015; Georges, Guillaume & Schiltz, 2020; Marlair, Crollen & Lochy, 2022). We propose that increasing variability, such as modeled most strongly by the 20 font stimulus set here, is therefore a promising avenue in future research targeting conceptual processing.

Using multiple fonts to increase variability has been used to advantage in some previous studies with Arabic numerals (Cohen, 2009; Vogel, Goffin & Ansari, 2015; Lochy & Schiltz, 2019; Guillaume et al., 2020; Finke et al., 2021; Marlair, Crollen & Lochy, 2022). Fonts have the advantage of likely retaining characteristic shape features so that they are easily and rapidly identifiable, although they are less variable than hand-drawn numerals as used here. Of course, variability can be supported through other means than diverse stimulus exemplars. Here, variability of image size was also applied at each stimulus presentation (Dzhelyova & Rossion, 2014); variability of stimulus position, orientation, contrast, etc. could also be used in future studies (e.g., as applied in some previous studies with Arabic numerals: Vogel, Goffin & Ansari, 2015; Guillaume et al., 2020; Finke et al., 2021).

Variability could also be reinforced through using larger stimulus sets, e.g., incorporating 0 and 1, and using multiple-digit numbers (although for parity, the ending Arabic numeral is inevitably limited; and even numerals ending in 0,2,4,6, and 8, may be inherently more closed and rounded than the odd numerals 1,3,5,7, and 9). Further, variability could be increased by testing stimuli in different stimulation modalities (e.g., Kiefer & Dehaene, 1997; Dickson et al., 2018; Finke et al., 2021) or stimulus formats. For example, studies on magnitude comparisons of near and far numbers have compared responses to numeral, word, and dot representations of numbers (Dehaene, 1996; Temple & Posner, 1998; Libertus, Woldorff & Brannon, 2007). While similarities in responses across formats suggest consistent conceptual processes, this approach is limited in that differences in responses across formats are difficult to attribute to stimulus properties or cognitive processing (see Libertus, Woldorff & Brannon, 2007; and Plodowski et al., 2003).

Another means to increase stimulus variability is to mix multiple formats of number representations, such as numerals, number words, dots, and finger signs (Marinova et al., 2021; Marlair et al., 2021; Marlair, Crollen & Lochy, 2022). However, physical stimulus confounds are as likely to combine across formats as to cancel out. For example, symbolic numerical representations are often built upon literal representations of the quantities they represent (Ifrah, 2000), and changing non-symbolic representations of quantity likely share similar confounds across stimulus formats. Moreover, investigations in some domains may best rely on a single representation of numbers, e.g., using only Arabic numerals that are familiar in the domain of mathematical equation processing. Overall, while increasing stimulus variability (through multiple means) is important, particularly for small stimulus sets, it may not be sufficient to overcome physical stimulus confounds.

### Responses to physical stimulus differences

It should be observed that physical stimulus differences greatly affected the EEG response amplitude. Significant asymmetry responses at 3.75 Hz were observed for all conditions of all stimulus sets, even those with high variability. These control asymmetry responses reflected contrasts of two group of four numerals, that cannot readily be explained conceptually: in both control conditions, the numeral sets were balanced for parity and approximately for magnitude.

Here, controls of non-conceptual numeral groups were used to estimate the impact of physical stimulus confounds on the parity response (as in Guillaume et al. 2020, except that in that study only half of the trial repetitions were tested for the control, which could account for the lower baseline-dependent control responses reported). Two control conditions were used here to give measures of predicted low and high visual differences across numeral groups. The Weak control condition was designed intuitively to have a large amount of shape differences across numerals groups: straight and open shapes (2,4,5,7; all containing at least one straight line segment, and only one of which contains a closed element) *vs.* curvy and closed shapes (3,6,8,9; none of which contains a straight line segment, and three of which have at least one closure). Not unrelatedly, the Weak control also had the largest differences across numeral groups in non-background image space, perimetric complexity, spatial frequency, and physical similarity (**Table S1**). The EEG response amplitude to the Weak control was about double that to the Strong control consistently across three stimulus sets, i.e., all but 20 drawn, and was much larger than the Parity response for the 1 font stimulus set (**Fig. 3**). Instead of being well-matched to the Parity condition, the Weak control served here as a dramatic demonstration of the impact of potential stimulus confounds.

This emphasizes the importance that studies address (intrinsic) physical stimulus confounds with controls, to limit the possibility that stimulus confounds, rather than conceptual representations, could account for reported effects. Importantly, this issue is not only relevant for non-symbolic numerical representations, but is demonstrated here for symbolic, Arabic numerals (see also Cohen, 2009). To be most informative, these controls should be as well-matched as possible: for example, while it is possible to use nonmeaningful characters, letters, foreign language letters, and/or symbols as a control condition (e.g., Iguchi & Hashimoto, 2000; Price & Ansari, 2011; Park et al., 2014; Fraga-Gonzalez et al., 2022), their visual attributes might not appropriately match those of experimental interest. For another example, a control condition of non-frequency-tagged trials, leading to no frequency-tagged responses, is not informative about potential physical stimulus confounds (Marinova et al., 2021).

With the two control conditions tested here, coarsely targeting shape, we are not able to point to which stimulus attributes most affected the neural responses. By testing additional control numeral groups, it may be possible to further probe which physical differences relate to larger EEG response amplitudes (e.g., cumulative perimeter, area, spatial frequency; for other approaches to identify visual features influencing non-symbolic representations of number with fMRI: Cohen Kadosh et al., 2007; Wilkey et al., 2017). We identified 15 possible control conditions that could be tested with the numerals 2-9 (each consisting of two sets of four numerals equated for parity and excluding large magnitude differences across sets). Understanding the impact of different physical features on responses with different methodologies would be highly informative for future studies’ design control. It could also be argued that testing further control conditions would increase the specificity of a higher response amplitude to a targeted conceptual condition.

### Qualitative response aspects

In theory, neural responses to numerals that reflect conceptual processing could be dissociated from responses merely driven by physical stimulus differences not only by amplitude, but by qualitative aspects, such as latency or localization. For example, it has been suggested that EEG responses reflecting conceptual processing of numerosity may occur later in time than those to low-level changes in visual shape of dot arrays (Soltesz & Szucs, 2014; Akbari et al. 2022); but this is contradicted in other studies reporting early processing of numerosity with EEG (Park et al., 2016; see also: Liu et al., 2018). Here, little temporal information could be extracted, given the high stimulation rate of 7.5 Hz, i.e., stimulus presentation every 133 ms, and asymmetry rate of 3.75 Hz (and low phase reliability: **Fig. S2b**).

An EEG scalp topography lateralized over the occipito-temporal/parietal cortex is generally interpreted as involving specialized numerical processing (e.g., Dehaene, 1996; Marinova et al., 2021; but also at medial-occipital sites: Pinel et al., 2001; Guillaume et al., 2018; George, Guillaume & Schiltz, 20202), in line with the parietal cortex being identified as highly relevant from lesion and neuroimaging studies, along with other cortical regions (reviews: Hubbard et al., 2005; Moeller, Willmes & Klein, 2015). However, differences in physical stimulus properties might account for extremely different response topographies targeting similar high-level processes (e.g., responses to dot quantities that are occipitotemporal: Hyde & Spelke, 2009; Marinova et al., 2021; or medial-occipital: Hyde & Spelke, 2009; Guillaume et al., 2020). On the other hand, similar topographies targeting conceptual and visual responses have also been reported (Van Rinsveld et al., 2020; but see Park et al., 2016).

Here, the asymmetry responses to parity at 3.75 Hz had lateralized occipitotemporal topographies; however, the control topographies within each stimulus set had similar topographies (as in Guillaume et al., 2020). Moreover, the responses to visual stimulation at 7.5 Hz were also lateralized over the occipitotemporal cortex (**Fig. S1**), suggesting that stimulus factors may contribute to lateralization as well as conceptual processing. One possibility is that shape processing, for Arabic numerals or for perceptually grouped small-number dot arrays, may invoke more visual processing beyond the early visual cortex (for less lateralized responses to larger dot numerosities: Libertus, Woldorff & Brannon, 2007; Hyde & Spelke, 2011; Guillaume et al., 2018 *vs.* Marinova et al., 2021; see also Han et al., 2002; Murray et al., 2002). This further suggests that in this context, topography lateralization is sensitive to physical stimulus confounds, rather than being a reliable indicator of conceptual responses.

There were pronounced differences in the side of lateralization here across stimulus sets: the response to Parity was more left-lateralized with 1 font but more right-lateralized with 20 drawn, and bilateral with 10 fonts and 10 mixed (**Fig. 3**). The control responses also varied somewhat within each stimulus set, in being lateralized more over one hemisphere or bilateral (**Fig. S1**). Response lateralization has been inconsistent across or even within studies on numerical cognition, with changes in numbers and stimulus properties (e.g., see Figs. 4 and 7 of Libertus, Woldorff & Brannon, 2007; Fig. 2 of Marlair, Crollen & Lochy, 2021), which is often attributed to conceptual differences. However, here, such effects depended on the stimulus set, even for the same numeral groupings, suggesting that physical stimulus factors may be sufficient to drive (the side of) EEG response lateralization, at least with relatively weak conceptual responses.

Although the group-level scalp topography for parity at 3.75 Hz did not appear to differ from that to the control conditions, the consistency of the response scalp topography across individual participants was strongest in the parity condition (**Fig. S2a**). Similarly, the phase of 3.75 Hz responses was also more consistent across individual participants (**Fig. S2b**), although phase is more reliable with higher amplitude. Future studies could investigate individual differences as a means of probing conceptual responses to numbers. Overall, a lack of qualitative distinction for parity *vs.* control responses here underlines the importance of controlling stimulus confounds in order to target conceptual responses to symbolic numbers.

### EEG frequency-tagging

Our paradigm benefited from the advantages of EEG frequency-tagging, including automaticity, specificity, consistency, and objectivity (in numerical cognition: Libertus, Brannon & Woldorff, 2011; Park, 2018; Guillaume et al., 2020; Van Rinsveld et al., 2020; Marinova et al., 2021; Marlair et al., 2021; Marlair, Crollen & Lochy, 2022). *Automatic* aspects of numerical processing, occurring without instruction or directed attention, can be measured because participants need not be assigned a (number-)related task. *Specificity* of numerical responses is protected from procedural variability in instruction and task performance, i.e., participants’ understanding of the language, mathematical terms, motivation, test anxiety, etc. Also, the absence of a related task enables the same paradigm to be used *consistently* with adults, children, infants, patient populations, and non-human species (e.g., Libertus, Brannon & Woldorff, 2011; Park, 2018). Frequency-tagging confers a high signal-to-noise ratio and *objectivity* in response identification and measurement (Regan, 1989; Rossion, 2014; Norcia et al., 2015). Moreover, thanks to frequency-tagging, the asymmetry responses at 3.75 Hz could also be contrasted to the general responses to visual stimulation at 7.5 Hz, which did not differ across conditions within each stimulus set.

## Conclusion

In conclusion, we suggest that neural responses may reflect conceptual numerical processing of Arabic numerals, however there is an important influence of physical stimulus confounds that should be taken into account in studies targeting conceptual responses.

## Acknowledgments

This work was supported by the SNAMath INTER project [INTER/FNRS/17/1178524 to CS] funded by the Luxembourgish Fund for Scientific Research (FNR, Luxembourg). We thank Vera Hilger and Brenda Gilson for their valuable assistance in piloting and data collection, and Bruno Rossion for providing helpful feedback on an earlier version of this manuscript.

