## Supplemental for "Identifying conceptual neural responses to symbolic numerals"

### Supplemental Material

#### Behavioral results

Performance at the fixation cross luminance change task (brief changes from dark gray to white) was performed with high accuracy (M = 96.4%; SE = 1.14%; range = 84.7% to 100%) and fast response times (RT; M = 476 ms; SE = 13.9 ms; range = 408 to 594 ms) overall, suggesting that participants paid attention to the screen and understood the task instructions. For both accuracy and RT, there was a main effect of Stimulus Set (accuracy: $F_{3,42} = 5.07$, $p = .004$, $\eta_p^2 = 0.27$; RT: $F_{1.8,25.4} = 44.1$, $p < .001$, $\eta_p^2 = 0.76$). There was not a main effect of Condition or an interaction (accuracy: F's < 2.2, p's > .14, $\eta_p^2$'s < 0.14; RT: F's < 0.97, p's > .41, $\eta_p^2$'s < 0.07). Performance was lowest for the 20 drawn stimulus set (accuracy: M = 90.0%; SE = 3.8%; RT: M = 552 ms; SE = 21.4 ms), with all other means over 98.0% for accuracy and ranging from 453 to 457 ms for RT. Worse performance with the 20 drawn stimulus set is likely due to the distracting complexity and color, including white, of this stimulus set.

| | **Parity** | **Strong control** | **Weak control** |
|---|---|---|---|
| **Non-background image space** | 1.35 | 1.67 | 2.20 |
| **Perimetric complexity** | 2.90 | 3.66 | 37.7 |
| **Image energy** | 0 | 0 | 1.5 |
| **High spatial frequency** | 0.024 | 0.25 | 0.65 |
| **Physical similarity** | 0.11 | 1.04 | 1.39 |

***Table S1. Image statistics for the 1 font stimulus set****: values represent the differences between numeral groups. Therefore, the higher the values here, the more that visual property might be expected to contribute to higher EEG asymmetry responses at 3.75 Hz. Non-background image space represents a measure of "retinal size". Image energy was defined as the lowest frequency (cycles/image) that contains 80% of image energy. High spatial frequency was defined as the* *percent of image energy above 10 cycles/image. Physical similarity is the ratio of overlapping to different line segments.*

| | **Parity** | **Strong control** | **Weak control** |
|---|---|---|---|
| **1 font** | 14.4 | 12.8 | 32.1 |
| **10 fonts** | 8.70 | 4.94 | 13.3 |
| **10 mixed** | 7.94 | 2.74 | 8.12 |
| **20 drawn** | 21.2 | 8.51 | 9.11 |

***Table S2. Z-scores for asymmetry responses****, harmonic-summed at the occipito-temporal ROI ($Z > 1.64$ corresponds to $p < .05$, 1-tailed).*

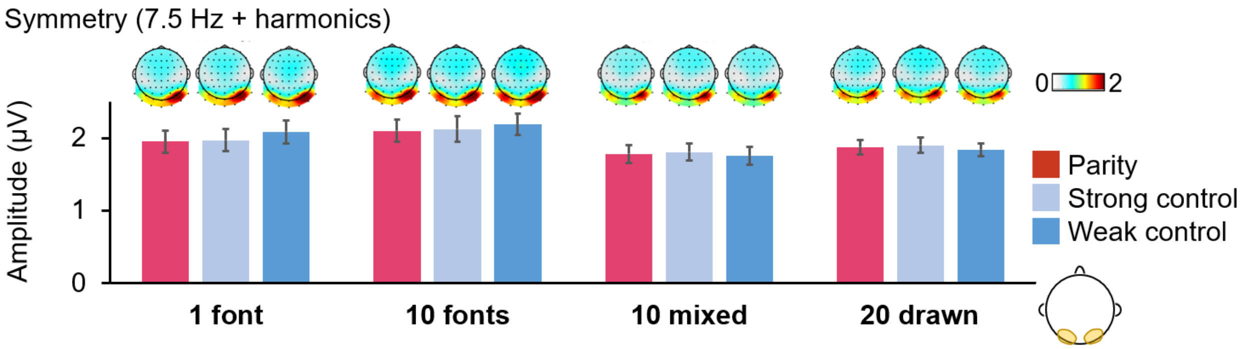


***Figure S1. Symmetry responses at 7.5 Hz and harmonics*** *(all notations as in Fig. 3).*

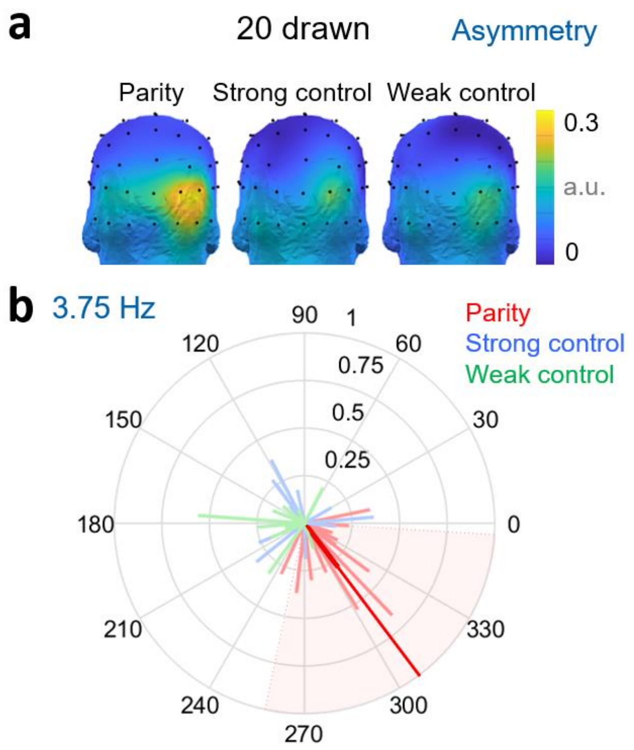


***Figure S2*. *Topography consistency and phase for the 20 drawn stimulus set. a)*** *Individuals' topographies were normalized before being averaged together: these head plots thus indicate the reliability of spatial response distributions across participants, rather than absolute amplitude differences.* ***b)*** *Vectors represent individual participant data, with the amplitude defining the length of the vector, and phase at 3.75 Hz its angle. Only the Parity condition was reliable enough across participants to indicate a shaded, 95% confidence interval of the mean (thick dark line, with extension to the edge of the graph for simpler value estimation).*
